# Cognitive penetrability of scene representations based on horizontal image disparities

**DOI:** 10.1101/2022.03.09.483682

**Authors:** Yulan D. Chen, Milena Kaestner, Anthony M. Norcia

**Author notes:** These authors contributed equally to the work (Yulan D. Chen, Milena Kaestner).

## Abstract

The structure of natural scenes is signaled by many visual cues. Principal amongst them are the binocular disparities created by the laterally separated viewpoints of the two eyes. Disparity cues are believed to be processed hierarchically, first in terms of local measurements of absolute disparity and second in terms of more global measurements of relative disparity that allow extraction of the depth structure of a scene. Psychophysical and oculomotor studies have suggested that relative disparities are particularly relevant to perception, whilst absolute disparities are not. Here, we compare neural responses to stimuli that isolate the absolute disparity cue with stimuli that contain additional relative disparity cues, using the high temporal resolution of EEG to determine the temporal order of absolute and relative disparity processing. By varying the observers’ task, we assess the extent to which each cue is cognitively penetrable. We find that absolute disparity is extracted before relative disparity, and that task effects arise only at or after the extraction of relative disparity. Our results indicate a hierarchy of disparity processing stages leading to the formation of a proto-object representation upon which higher cognitive processes can act.

## Introduction

Surfaces and objects in natural scenes can be distinguished based on gradients and discontinuities in a wide range of local features and the coding of gradients and discontinuities involves non-local feature measurements. The earliest stages of so-called figure-ground segmentation have been extensively studied for motion ^1-6^ and orientation discontinuities ^7-11^ in V1 and V2 of macaque where sensitivity to discontinuities in these features is present.

Discontinuities in binocular disparity also provide robust signals for figure ground segmentation and can do so in the absence of any other cues ^12^. Binocular disparity is an interesting case, because unlike the case of motion and orientation discontinuities, were both local cues themselves and discontinuities in these cues are robustly signaled in V1, only local disparities – so-called absolute disparities – are robustly represented in V1, with the representation of disparity discontinuities (relative disparities) being deferred until V2 ^13-16^. Importantly, cells in V1 are tuned for the absolute disparity of both correlated and anti-correlated stereograms ^17^. Only the former support a percept of depth. Thus, the neural basis of perceptual depth must lie outside of V1 ^18^.

In a parallel line of research, the human psychophysical and oculomotor literatures have also suggested that absolute disparity representations are perceptually inaccessible. When confronted with a large, uniform random-dot stereogram whose absolute disparity is slowly changed over time, observers are unaware of the motion in depth ^19,20^, even under stabilized image conditions that produced large absolute disparities that could not be compensated by vergence eye movements ^21^. The addition of a disparity reference, creating relative disparity cues, greatly increased the percept of motion-in-depth. Stimuli containing vertical absolute disparity also evoke no percept of motion-in-depth, but can drive vergence ^22^. A comparison of the relationships between absolute and relative disparity thresholds and vergence noise has suggested that absolute disparity, *per se* is perceptually inaccessible, something the authors termed the absolute disparity anomaly ^23^. The geometric definitions of absolute and relative disparities are illustrated in Figure 1a.

**Figure 1:**
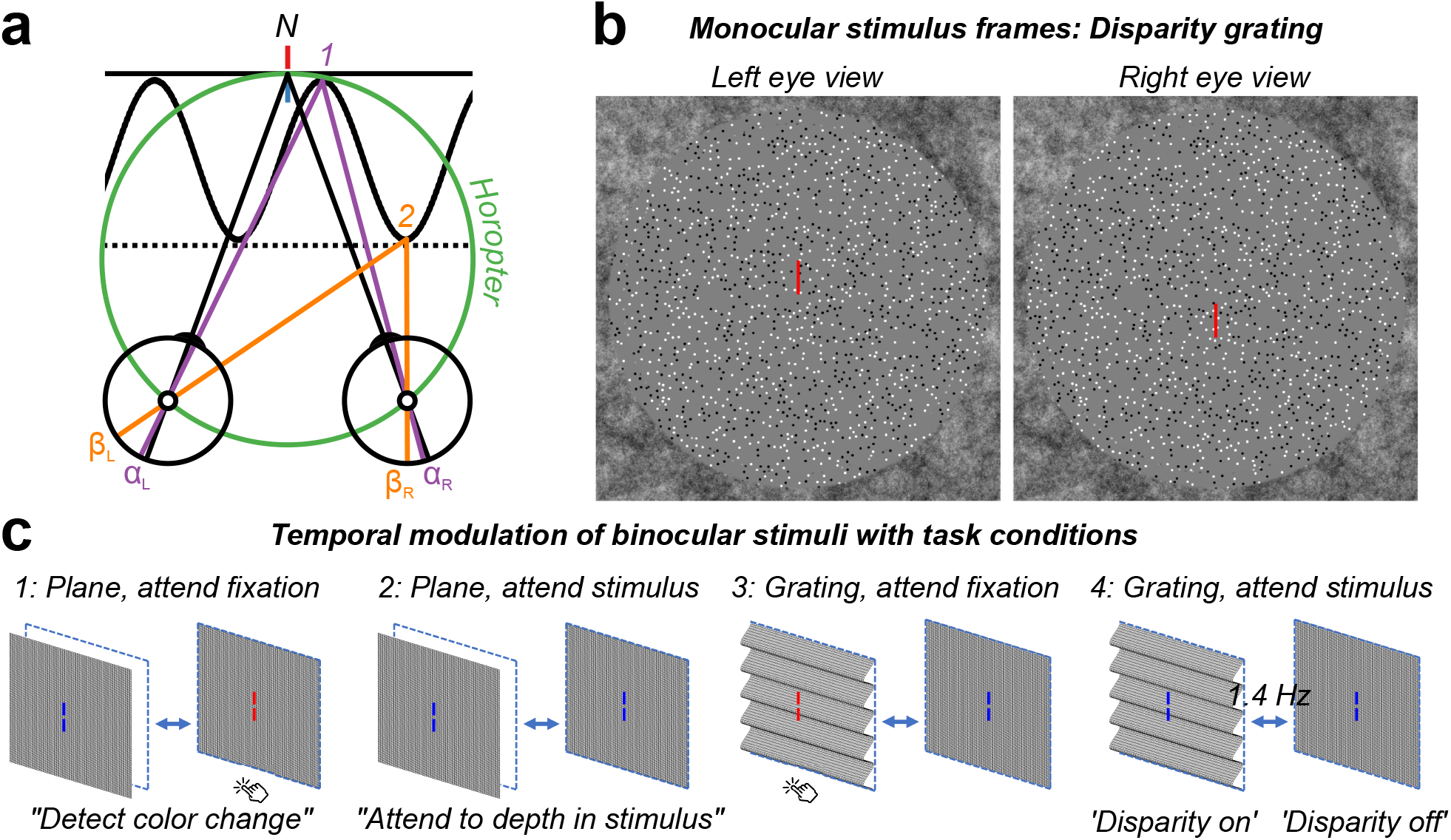
**a:** Top-down schematic illustration of absolute and relative disparity. Fixation point N (nonius lines) is on the zero-disparity plane defined by the horopter (green Vieth-Muller circle). Point 1 (purple) is also on the horopter and angles α_L_ and α_R_ are equal, meaning that the absolute disparity given by α_L_ − α_R_ is zero. Point 2 (orange) is either on a second plane (dotted line) or on the peak of a disparity grating (sinusoidal line). Here the absolute disparity, β_L_ − β_R_, is non-zero. The relative disparity is the difference between the two absolute disparities, (α_L_ − α_R_) − (β_L_ − β_R_), and varies depending on the location on the sinewave but its magnitude is independent of fixation. Disparity along the second plane (dotted line) is constant at a non-zero value. **b:** Random-dot stereopair used in the main experiment depicting a sinusoidal disparity grating (crossed disparity when cross-fused). The dots in the actual experiment were dynamic. **c:** Schematic of different stimuli, task conditions, and temporal modulation between crossed disparity, ‘disparity on’, and zero disparity, ‘disparity off’ phases. Participants were asked to detect a brief color change on the fixation lines, or to attend to the depth modulation of the stimulus, for either plane (absolute disparity only) stimuli or grating (absolute and relative disparity) stimuli. To generate a visual-evoked potential, all stimuli modulated between disparate and non-disparate states at 1.4 Hz.

If absolute disparities are perceptually inaccessible, their processing should be relatively immune to the observer’s task. By contrast, responses to perceptually relevant relative disparities may be subject to task effects. The literature on orientation-based figure-ground processing has posited a significant role for top-down inputs to the figure-ground process and further suggests a time-course for attentional modulation relative to the extraction of local features and local feature relationships. V1 cell responses of macaque in an orientation-defined form task show that the effect of task occurs after orientation discontinuities and by implication orientation tuning *per se* was established ^24^. Later work ^25^ suggested a model comprised of an initial feature registration stage, here local orientation, that was followed by detection of the feature-difference boundary, which was then followed by attention-dependent enhancement of the figure region and later suppression of the background. In related work ^26^, border-ownership relations in a display involving separated or occluding figures showed that some cells signaled border ownership independent of attention, but that in a substantial number attention acted after border ownership was established in the case of separated figures and at the same time as border ownership was established of occluded figures.

Task effects on neural responses to absolute and relative disparity cues have not been studied before, but there are clear predictions for what should be observed from the existing literature where absolute disparity processing should be relatively immune from the effects of task. The extent to which relative disparity coding is influenced by task is a more open question. Here, we compared 128-channel evoked Visual Evoked Potentials (VEPs) to dynamic random-dot stereograms (DRDS) that either portrayed a flat surface moving in depth (changing absolute disparity) or a flat surface changing to a corrugated one (changing absolute and relative disparity). For readers who can free-fuse stereograms, a crossed-disparity (static) random-dot stereogram depicting a disparity grating is shown in Figure 1b. The use of DRDS stimuli allowed us to isolate disparity-related responses from responses due to monocular stimulation, and the use of the grating vs. plane stimuli allowed us to manipulate the availability of relative disparity information. To modulate potential top-down influences on these responses, we recorded under task conditions in which the disparity modulation was either task relevant (‘attend stimulus task’) or task-irrelevant (‘attend fixation task). Schematic illustrations of stimuli, temporal dynamics, and task conditions are illustrated in Figure 1c.

We find that the initial evoked response to disparity change is the same for the disparity plane and disparity grating stimuli, suggesting that the leading edge of the response reflects the processing of absolute disparity information that is common to the two stimulus conditions. We also find that the response to the disparity grating condition is more affected by task, and that this task modulation occurs at or after the extraction of relative disparity.

## Results

### Topography of disparity responses

We used a spatial filtering approach (Reliable Components Analysis, RCA; see Methods: VEP Signal Processing and Reliable Components Analysis, also Figure 7b) to reveal clusters of electrodes that responded in a systematic manner across stimulus presentation trials ^27^. RCA improves SNR and reduces the dimensionality of our 128-channel electrode data into a smaller number of components, whilst also revealing underlying, physiologically plausible neural sources that are illustrated in Figure 2. Here, we show the topographies of the first reliable component, RC1, highlighting a cluster of midline occipital electrodes that respond reliably across trials of the plane (a) or the grating (b) stimulus. Sensor-space weights were learned on ‘nF1 clean’ filtered data in the time domain (see Figure 7a(iii)), collapsing across task conditions (‘attend fixation’ and ‘attend stimulus’) but between disparity stimulus types (plane vs. grating). The resulting topographies for the first component RC1, shown in Figure, 2 were produced via a forward model generated from the learned weight vectors ^27,28^ and are visually similar, and consistent with neural sources in V1, V2, V3 and V3A ^29-32^. We note that the response to the grating is somewhat more diffuse, commensurate with a more widespread neural response to relative disparities. In the remainder, we present results from these topographies, where 128-channel data from each condition were weighted by either the ‘plane’ or the ‘grating’ RC1 vector to generate group-level waveforms and error estimates.

**Figure 2:**
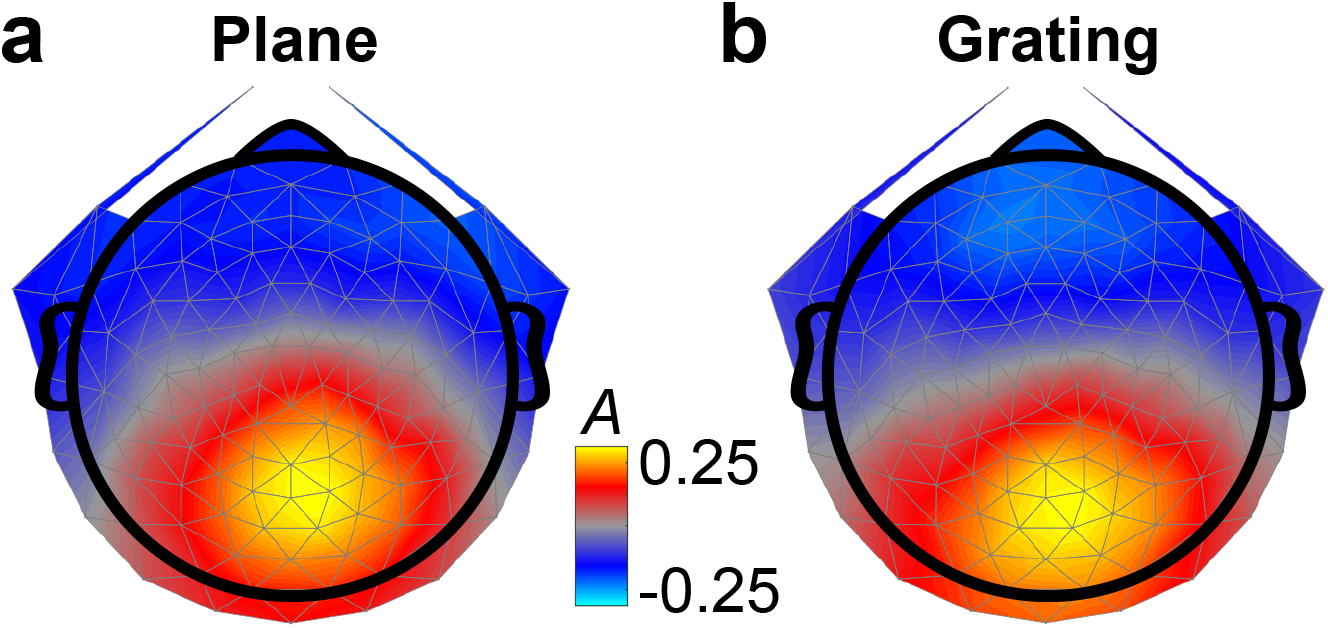
Topographies of the first reliable component for responses to plane and grating stimuli (N=20). Units of the forward model (A) were maximal over midline occipital electrodes, indicating that neurons in early visual cortex were responding systematically across trials. Small differences between the plane (**a**) and the grating (**b**) responses include that the grating response is shifted more towards posterior electrodes and is broader.

### Time-course of disparity plane and grating responses

To assess the relative timing of plane and grating disparity processing, we first compare responses of the disparity plane to the disparity grating during the nonius color-change task (Figure 3a). The disparity-evoked responses both show a positive peak at ∼100 ms post-disparity onset and diverge significantly from one another at 172 ms on the downward slope of a negative-going potential. The initial portion of the response (∼100 – 172 ms) is thus the same to within measurement error between the conditions, with the negative deflection being larger for the relative disparity condition. Notably, the response to the disparity grating is sustained for the duration of the disparity-on phase, whilst the response to the disparity plane is transient at the disparity-on and disparity-off transitions. Differences in response profile drive the long period of significant differences between the waveforms post 172 ms.

**Figure 3:**
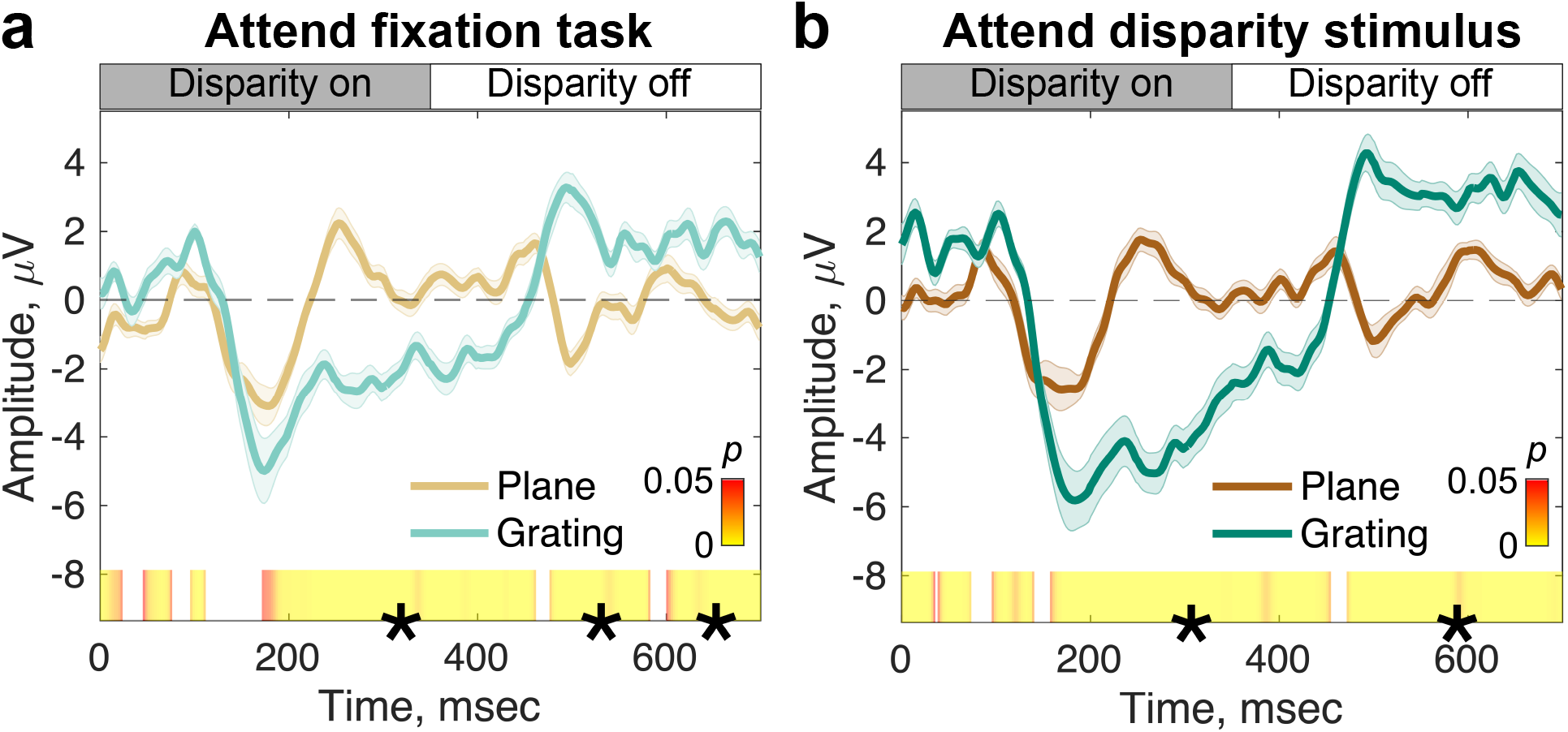
Comparisons of EEG responses to disparity changes for plane and grating stimuli under ‘attend fixation task’ (**a**) and ‘attend disparity stimulus’ (**b**) task conditions (N=20). In both panels, EEG waveforms are reconstructed via the nF1 clean filter and are weighted by the first reliable component (topographies in Figure 2). Waveforms are group-level mean responses to a single stimulus cycle consisting of one ‘disparity on’ and ‘disparity off’ transition. Errors are ±1 SEM. The heat bar indicates the p-value significance of the plane (brown colors) vs. grating (green colors) responses calculated via permutation-based paired T-tests. Segments marked with (*) indicate sequential, significant time points that pass the threshold for run-length correction.

We also compared responses to the two disparity types under ‘attend stimulus’ conditions (Figure 3b). Here the disparity-evoked responses also show positive peaks at ∼100 ms, but they diverge at 158 ms rather than at 172 ms in the attend fixation condition. Asking the observers to attend to the changing disparity stimulus rather than to the nonius lines can thus speed up the differentiation of the plane stimulus that contains only absolute disparity, from the grating stimulus which contains both absolute and relative disparity, but not to the extent that it eliminates the common initial positive/negative deflections.

### Does task have equivalent effects on plane and grating responses?

We asked whether the plane and grating conditions are equally affected by task. As shown in Figure 4, responses to the plane condition under the two tasks are similar, overlapping entirely during the two transient phases of the response that occur at disparity-on and disparity-off transitions (negative peaks at 176 ms and 498 ms for both task conditions). The brief periods of significance at the start and end of the cycle are likely driven by residual wrap-around effects of later activity (for further discussion see below) and occur before and after stimulus-evoked activity associated with the disparity change.

**Figure 4:**
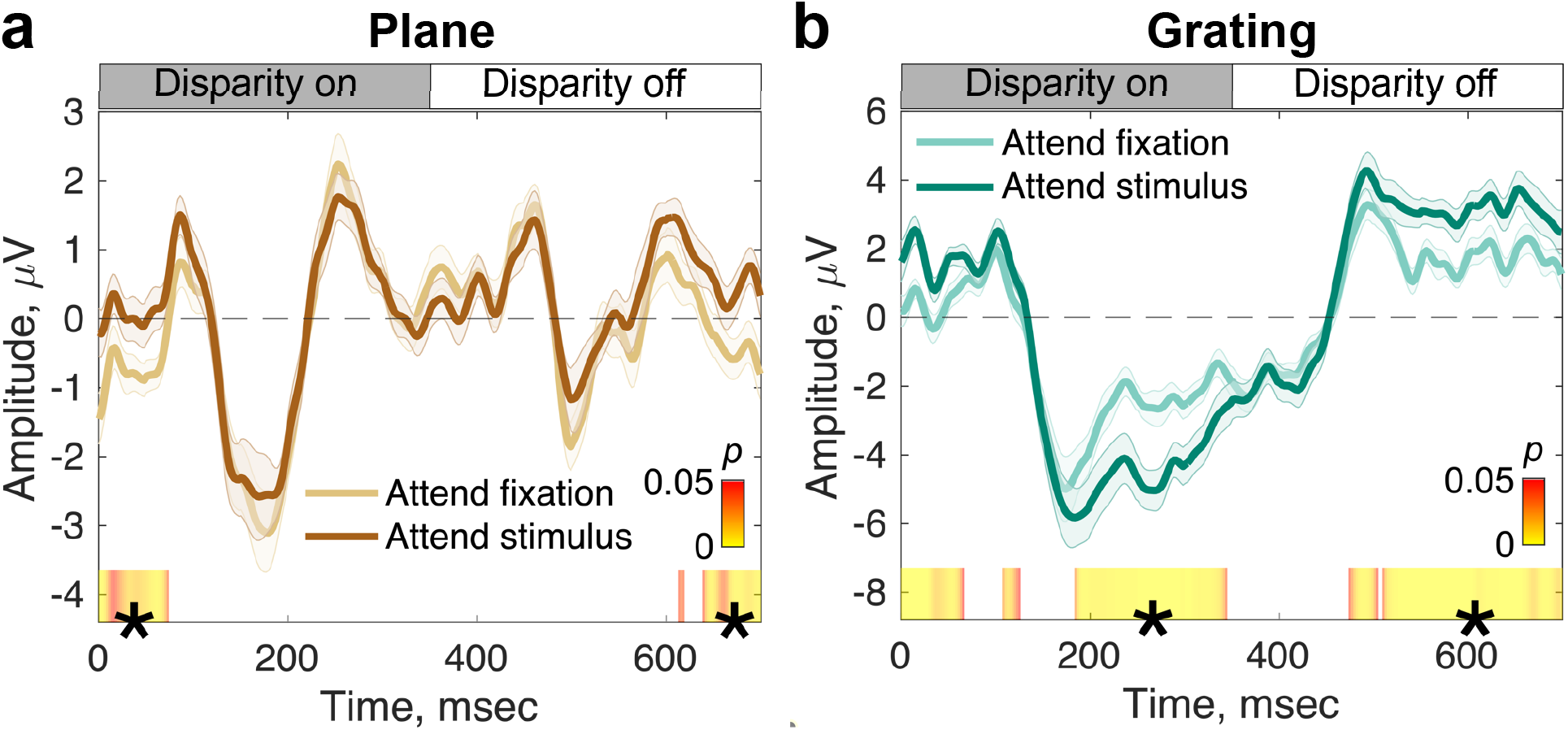
EEG responses to plane and grating disparity stimuli under different task demands (N=20). **a:** Responses to the plane (pure absolute disparity) stimulus. Responses during attend fixation (light brown) and attend stimulus (dark brown) conditions are highly similar and transient waveforms are observed at both disparity on and disparity off stimulus transitions. **b:** Responses to the grating (relative and local absolute disparity) stimulus. Responses during the attend fixation (light green) and attend stimulus (dark green) are initially similar up to ∼184 ms when they begin to diverge. For both panels, data are the group mean waveform in RC1, and errors are ±1 SEM. Data have been filtered using the nF1 clean filter. Heat bars indicate significance (paired-samples T-test) and segments marked with (*) indicate sequential, significant time points that pass the threshold for run-length correction.

Grating stimuli, as opposed to plane stimuli, contain more perceptually relevant information with the addition of relative disparity cues, and thus we expected to find greater effects of task here. As was the case for the plane disparity conditions, the leading edge of the responses for the grating stimuli are identical (102-184 ms), but the ‘attend stimulus’ condition results in a larger negative-going deflection that is sustained throughout the ‘disparity on’ phase of the stimulus (Figure 4b). The magnitude of the response significantly differentiates the two task conditions, starting at 184 ms after disparity onset and lasting through 344 ms. This effect occurred after the differentiation of the plane and grating responses at 158 or 172 ms, depending on attentional demands (Figure 3). Task effects are also seen at comparable latencies after disparity offset. Although our task manipulation resulted in large signal deflections later in the stimulus cycle, it did not affect the initial phase of the response at disparity-onset.

### Separating transient and sustained components via spectral filtering

‘Odd’ and ‘even’ filters in the Fourier domain can be used to reveal transient and sustained response components that have been associated with the extraction of plane and grating disparity signals respectively (Kaestner et al., in revision; McKeefry et al., 1996). The transient response component to the grating stimulus was separated by reconstructing the waveform using only even harmonic response components (2F1, 4F1, 6F1, etc.), and the sustained component using only odd harmonic components (1F1, 3F1, 5F1, etc.). Filtering further highlights the differences in the response profiles for the grating and the plane conditions, where the response to the plane is dominated by even harmonics (Figure 5b) whilst the response to the grating is contains both odd (Figure 5c) and even (Figure 5d) harmonic responses. For the grating stimuli, odd harmonic responses are at least a factor of two larger in amplitude than even harmonic responses. Thus, the plane stimulus, containing only absolute disparities, drives an even symmetric response with transient on and off responses, whilst the grating stimulus, containing both absolute and relative disparities, drives a response with both odd and even harmonic components. This pattern of results associates the odd harmonics with the processing of relative disparity.

**Figure 5:**
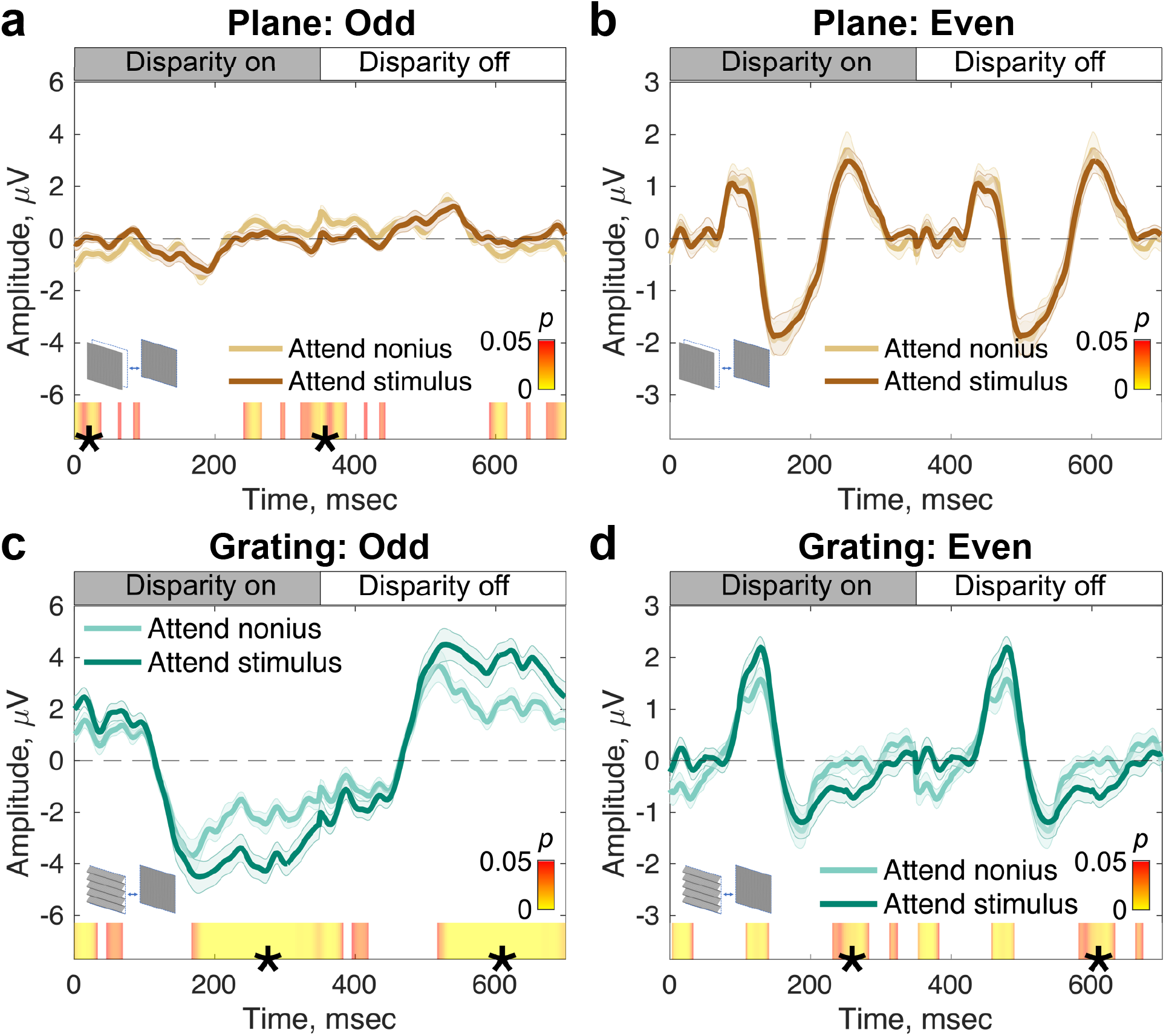
Task effects on EEG responses to disparity stimuli under different Fourier filtering regimes that highlight sustained vs. transient response components (N=20). **a** and **b** show responses to the plane stimulus (absolute disparity); **c** and **d** show responses to the grating stimulus (relative and local absolute disparities). The “Odd” filter (left column) preserves the odd harmonics of the fundamental frequency (1F1, 3F1, 5F1, etc.) and highlights sustained response components driven by the amplitude of the 1F1 response. Where this signal is present (**c**), there is a significant effect of task. The “Even” filter (right column) preserves the even harmonics of the fundamental frequency (2F1, 4F1, 6F1, etc.) and reveals transient, temporally symmetric responses at disparity on- and off-sets. There is no task effect for the plane stimulus (**b**) and there are some small effects for the grating (**d**), but these are localized outside the main positive/negative deflections that demarcate the stimulus transitions. Data are group-level averages weighted by RC1, errors are ±1 SEM and the color bars indicate significance (paired-samples T-test). Segments marked with (*) indicate sequential, significant time points that pass the threshold for run-length correction.

How does the filtering approach reveal effects of task? For the plane stimulus, the even-filtered response shows no measurable effect of task and a small odd-harmonic response with task effects at the edges of the cycle average (Figures 5a and 5b). The odd harmonic response in the grating condition is by contrast ∼8 times larger than in the plane condition (compare Figures 5a and 5c) and shows substantial effects of tasks, starting at 167 ms (Figure 5c) – 17 ms earlier than in the unfiltered data. Like in the case of the unfiltered data, there is an initial period of response that is not modulated by task between 100 and 167 ms. The even-filtered response to the grating condition (Figure 5d) has measurable effects of task at 230 ms after a period between 75 and 200 ms where the disparity-specific response is present but not affected by task. The onset of the earliest disparity specific response is now more clearly defined in the even filtered responses, and occurs at ∼75 ms for both the disparity plane and grating conditions.

## Discussion

By using a combination of high-density EEG, a spatio-temporal decomposition approach, and behavioral task manipulations we have mapped out the temporal order of the early stages of disparity processing and their modulation by task. We found that during the earliest time-points after disparity onset, responses to the plane and grating conditions, and by implication responses to absolute and relative disparity information, did not differ from each other and were not dependent on task. We also observed a brief period where relative disparity information was registered independent of task after which the effect of task manifested primarily on responses we associate with the extraction of relative disparity. In the following we detail the timeline of disparity processing with respect to our own data and to the broader context models of attentional influences on scene processing.

### Time-courses for the onset of absolute and relative disparity processing

DRDS stimulation isolates disparity-specific responses as the stimuli contain no monocular cues for the disparity changes that elicit the evoked response. By varying the spatial structure of our stimuli, we could compare responses to a stimulus that contained only absolute disparity (the plane condition) to one that also contained relative disparity (the grating condition). Our spectral filtering approach separated time-courses that largely split the response into relative-disparity sensitive (odd harmonics) and relative disparity insensitive (even harmonics) response components. The even harmonic waveforms show an onset-time of ∼75 msec for both plane and grating stimuli, while the robust odd-harmonic waveform derived from the grating response had an onset time of ∼100 msec, *e*.*g*. ∼25 msec later. Thus, the positive peaks at ∼100 msec in the full waveforms in Figure 3 come from the even-harmonic, transient response component. Previous VEP studies have uniformly reported the presence of a negative going component after disparity onset or change ^33-35^. The positive peak that we see here at ∼ 100 ms has been only infrequently reported (Skrandies, 2001). This peak is small and may thus have been difficult to record with conventional signal averaging and low-channel count recording approaches. Our filtering approach thus provides robust access to this earliest stage of disparity processing, situating the onset of disparity responses to be within ∼25 msec of the onset of cortical response to image contrast (see ^36^ for detailed discussion).

The initial stage of disparity processing appears to be largely unaffected by the presence of relative disparities the display. Responses to the disparity grating stimulus which contains both absolute and relative disparities diverge from the response to the plane stimulus (absolute disparity only) at 158 to 172 msec, *e*.*g*. well after the initial disparity-specific response occurs. This pattern of delayed relative disparity selectivity is consistent with what is known from the functional anatomy of single-unit recordings in macaque. Relative disparities are extracted in macaque V2, V3/V3A, V4 and IT, but not in V1 ^13,14,37-39^. Human fMRI results also suggest that relative disparity begins to be extracted in V2 and V3, but not in V1^32^. One thus expects EEG responses driven by relative disparity to be temporally delayed with respect to responses driven by absolute disparity. The previous single-unit studies have focused on tuning for relative disparity and have not presented time courses for the onset of relative *vs* absolute disparity selectivity as we do here.

By filtering the raw responses into even and odd harmonic components, we see that the odd-harmonic filtered component is strongly selective for relative disparity present in the grating condition. Moreover, the odd harmonic component is delayed relative to the even component, consistent with the former being driven by the presence of relative disparity in the grating stimulus. The 25 msec difference in odd-*v*s even-harmonic response onset (compare between Figures 5c and 5d) is thus consistent with the even harmonic response reflecting the processing absolute disparity which is present in both plane and grating stimuli and the odd harmonics reflecting the extraction of relative disparity which is unique to the grating stimulus.

Prior work has also indicated that odd-harmonic responses are likely to arise from mechanisms that are sensitive to relative disparity. Odd-harmonic responses are strongly dependent on the availability of multiple disparities in the visual field ^40-42^ consistent with the differences we see between plane and grating responses. Odd harmonic responses are tuned for cyclopean spatial frequency while even harmonics are not (Kaestner et al., in revision), consistent with the even harmonics arising from mechanisms sensitive to absolute disparity. Here we add to these results by demonstrating that relative disparity is extracted after an initial transient response to absolute disparity that can be linked to even harmonic response components.

### Effects of task on disparity-specific responses

The response to the plane condition is dominated by even harmonic response components for which no modulation by task was seen. Because the stimulus isolates absolute disparity, the lack of a task effect is consistent with absolute disparity processing being task independent. What about the case of even harmonic/transient responses, more broadly? The grating stimulus has both absolute and relative disparity information and the response to the grating condition also contains even harmonics. Here the response is largely, but not entirely devoid of task effects. A small effect of task is measurable at around 230 msec after a disparity change. This modulation is small and occurs well after the larger, sustained task effect manifests in the odd-harmonic response.

Measurable odd-harmonic activity is present in the plane condition which is ∼ 8 times smaller than what is recorded in the grating condition. Odd harmonic responses in the plane condition could arise from residual responses to incomplete isolation of relative disparity information from our display, from asymmetries in the response to direction of motion in depth or from asymmetries in the population response to zero and non-zero disparities. Some of this activity may be residual stimulus artefact that was not removed by the ICA filtering. The task effects that occur are at the times of disparity change and may therefore be due to wrap-around effects.

The largest and most striking task effects we measure arise for the grating stimulus and are brought to prominence through the ‘odd’ filter that emphasizes the VEP’s sustained response profile. We consistently found that the amplitude of the sustained negative-going potential was amplified when participants were attending to the stimulus directly. Thus, the observer’s task boosts neural responses to the relative disparity content of the grating stimulus, presumably via feedback from higher cortical areas. This modulation occurs at time points after the initial disparity encoding phases, from 167 ms onwards. The initial phase, from 75 – 158 ms is shared between grating and plane conditions, and is related to the initial extraction of absolute disparities. The period between 158-167 ms represents relative disparity coding that is pre-attentive. It is only after relative disparity has been extracted that the behavioral task is able to modify the neural response. Note that this time point may be too late to impact the transient responses seen either for the disparity plane, or for the even-filtered grating response. Alternatively, feedback connections may not target the anatomical substrate that generates the response to absolute disparity. These two explanations need not be mutually exclusive.

### Cognitive penetrability of disparity processing

Our data speak to a conceptual framework that is prominent in the attention literature – the notion of cognitive penetrability. Cognitive penetrability and its converse, cognitive impenetrability, relate to the question of how early in the visual pathway the effects of the observer’s task manifest. A substantial body of evidence, largely derived from the ERP literature, has suggested that the earliest stage of cortical processing is not subject to task effects. This issue has been studied by an examination of the leading edge of the evoked response to the onset of an image which manifests as a component (C1) that occurs over ∼50-80 msec. Image onset creates contrast, proto-object (see following section in Discussion), and object-level transients, but the leading edge reflects the initial extraction of contrast. While some studies have reported task effects on C1 ^43-45^, other studies have indicated that C1 is not cognitively penetrable (see ^46,47^ for reviews).

Here we also see an early response component – a positivity that arises at ∼75 msec after disparity onset that too is cognitively impenetrable. This activity is seen in both plane and grating conditions and is associated with even response harmonics that show only small effects of task a much later time-points.

### The role of “proto-objects” in attention

The processing sequence we observe suggests the following sequence of coding stages: two task-independent coding stages, the first starting at ∼75 msec involving the registration of absolute disparity which is followed by a brief period of task-independent coding of relative disparity. Task effects manifest after this point, but primarily in the odd-harmonic component of the disparity grating stimulus. This sequence of processing resembles the set of stages posited by psychophysicists from behavioral experiments ^48-50^. Within this framework, the initial processing of a scene involves the rapid extraction of luminance, contrast, and edge information, as well as three dimensional surface orientation, and groupings of related edge fragments, resulting in the formation of a “proto-object” representation that does not depend on attention ^48^, which is consistent with data from macaque V1 showing that enhancement of cell responses by image segmentation cues such as orientation contrast occurs in the absence of focused attention ^11,24,51^. A similar pattern has also been observed in an Event-Related Potential (ERP) experiment on texture segmentation ^52^.

A proto-object representation only loosely corresponds to everyday, recognizable objects and surfaces, but it goes considerably beyond coding of raw image statistics such as local orientation ^49^ and here, absolute disparity. Attention, on this view, acts only after the formation of the proto-object representation. Relative disparity information fulfills the criteria required as the basis of a proto-object representation, as it can readily support the segmentation of figure and background and the computation of 3D surface orientation. Moreover, a relative disparity representation is largely independent of fixation distance, providing a degree invariance in terms of form-from-disparity that would be useful for downstream invariant representations.

In our hands, the observer’s task modulates the response only once a relative disparity, *e*.*g*. a proto-object representation, has been established. In the case of texture-defined from processing in macaque V1, the representation of boundaries and surfaces defined by an orientation discontinuity were seen to be an automatic process, with attention acting only after these to figure/group processes had been established ^24^. Effects of task in an earlier study were also only seen at late time points, well after segmentation-dependent responses occurred ^11^. Our results for absolute and relative disparity also suggest a hierarchy of proto-object representations that transitions from being task-independent to task-dependent.

Task and relative disparity extraction also appear to interact in the opposite direction – task can influence the speed of relative disparity processing (see Figure 3). This two-way interaction between attention and proto-object formation has been previously observed in the case of the coding of border-ownership in macaque V2 ^26^. Differential cell responses to figure and background regions (*e*.*g*. the border ownership signal) manifested this distinction more rapidly when attention was directed to the cell’s preferred vs non-preferred stimulus, indicating an interaction between task and the extraction of a proto-object representation.

Other work on human disparity processing is consistent with the multi-stage model of proto-object formation and attention. By recording differences in ERPs between flat, zero disparity images and 3D images, the presence of disparity was found to modulate the response starting at 100 msec, but it was not until 150 msec that the magnitude of the difference potential predicted behavioral reaction time ^36^. The corresponding value for the onset of disparity sensitivity in the current study is ∼75 msec, and the onset of a task effect is 165-184. Importantly, both studies have found a relatively long period when a disparity-specific response has begun but is not modulated by task or task performance.

Disparity is the primary cue for depth-based image segmentation, but monocular occlusion cues also serve as depth segmentation cues. Prior work with occlusion as a depth cue has suggested that attention operates after depth-order information has been extracted, *e*.*g*. after proto-object formation ^53^. In that study, a set of horizontal bars moved at one temporal frequency and an orthogonally oriented set of bars moved at a different temporal frequency. Depth order was established by occlusion cues supported by color differences between the bars signaling which set was seen in as in front of the other. Observers were tasked with attending to vertically oriented bars that were seen as either in front or behind the horizontal bars depending on the occlusion cue. A higher harmonic response component of the response (4F) was independent of task, but a lower frequency component (2F) was. Importantly, only the lower frequency component signaled depth order. As the higher frequency components are likely to come from the leading edge of the response, the pattern of results is consistent with attention acting at a later stage of processing after depth order and thus scene segmentation rather than motion, *per se* had been extracted.

## Conclusion

By using stimuli that have a hierarchy of disparity cues, from locally accessible absolute disparities to non-local relative disparity, we have demonstrated a hierarchy of processing leading to the formation of a proto-object representation. This representation is based on relative disparity and is cognitively penetrable.

## Methods

### Participants

24 participants (14 female, mean age 30 years) were recruited from the Stanford community. They all had normal or correct-to-normal visual acuity with no history of ocular diseases and neurological conditions. Visual acuity was assessed with a Bailey-Lovie LogMAR chart (Precision Vision, Woodstock, IL) with acuity of 0.1 LogMAR or better in each eye, and less than 0.3 LogMAR acuity difference between the eyes. The RANDOT stereoacuity test (Stereo Optical Company, Inc., Chicago, IL) was used to test stereo-acuity with a passing score of at least 50 arc seconds. Four participants were excluded from analysis, three due to low dot update responses (see below, Methods: Stimuli and Methods: VEP Signal Processing and Reliable Components Analysis) and one due to technical issues during recording. Data from 20 participants were retained for analysis. Participants’ informed written and verbal consent was obtained before experimentation under a protocol approved by the Institutional Review Board of Stanford University.

### Visual display

Dynamic random-dot stereograms (DRDS) were displayed on a SeeFront 32” autostereoscopic 3D monitor with a TFT LCD panel display in which a lenticular lens system presents separate image data to the left and right eyes. The SeeFront monitors participant position in front of the screen with an integrated pupil location tracker, and the eyes’ separate images are merged via line-interleaving in real time to form a single 3D image from the viewpoint of the participant. Viewing distance from the screen was 70 cm, within the optimal range as per the manufacturer for typical adult inter-pupillary distances. The screen has a native resolution of 3840 × 3160 pixels (1920 × 1080 effective binocular resolution) and was refreshed at 60 Hz. Mean luminance was 50 cd/m^2^.

### Stimuli

DRDS stimuli were like those described in detail elsewhere (Kaestner et al., in revision). An example of side-by-side binocular half-images is shown in Figure 1B. In brief, circular DRDS half-images (radius = 13.65°) were constructed of non-overlapping, 100% contrast black and white dots (15 dots/degree^2^, dot size 6 arcmin diameter) on a mean gray background. Dot positions were pseudorandom and regenerated in new locations at 20 Hz (every 3^rd^ video frame). DRDS stimuli alternated at a rate of 2 Hz between ‘disparity on’ and ‘disparity off’ states (modulation depicted in Figure 1c). During the ‘disparity on’ phase, stimuli generated a percept of a flat plane at 6 arcmin crossed disparity (‘plane stimulus’), or a percept of a horizontally-oriented, sinusoidal disparity grating (0.5 cpd) where binocular dot pairs were shifted systematically from being perfectly correlated (0 arcmin disparity) at the trough of the disparity grating, to being horizontally offset to generate 6 arcmin disparity at the peaks of the grating (see also Figure 1a for a depiction of absolute disparities in the plane stimulus in both ‘disparity off’ and ‘disparity on’ states – points 1 and 2 – and relative disparities in the grating stimulus – differences between points 1 and 2). The plane stimulus contained only absolute disparities, whilst the grating stimulus contained both local absolute and global relative disparities that defined the sinusoidal grating. During the ‘disparity off’ phase of the stimulus cycle, dot pairs were perfectly correlated between the eyes, generating a percept of a flat plane at zero disparity. Thus, our DRDS stimuli generated Visual Evoked Potentials at two fundamental frequencies, F1 and F2, as well as their harmonics, *n*F1 and *n*F2. *n*F1 were multiples of the 1.2 Hz disparity change, and *n*F2 were multiples of the 20 Hz dot update response. These stimulus-driven, frequency-tagged EEG responses were useful in later filtering stages (see Methods: VEP Signal Processing and Reliable Components Analysis).

Figure 6 shows a schematic view of key elements in the stimuli. The DRDS was windowed using a 1/f fusion lock (39.5° square) presented in the periphery and at zero disparity, and which was static throughout each stimulus presentation and stabilized vergence eye position (Figure 6, ‘a’). There was a band of uncorrelated dots between the fusion lock and the DRDS stimulus (1.2° wide; Figure 6, ‘c’) which was sufficient to disrupt edge and binocular reference effects between the fusion lock and the disparity stimulus ^54^. Centrally placed nonius lines (Figure 6, ‘b’) were vertically separated and thus conveyed no horizontal disparity information, and together covered 1° of the stimulus vertically.

**Figure 6:**
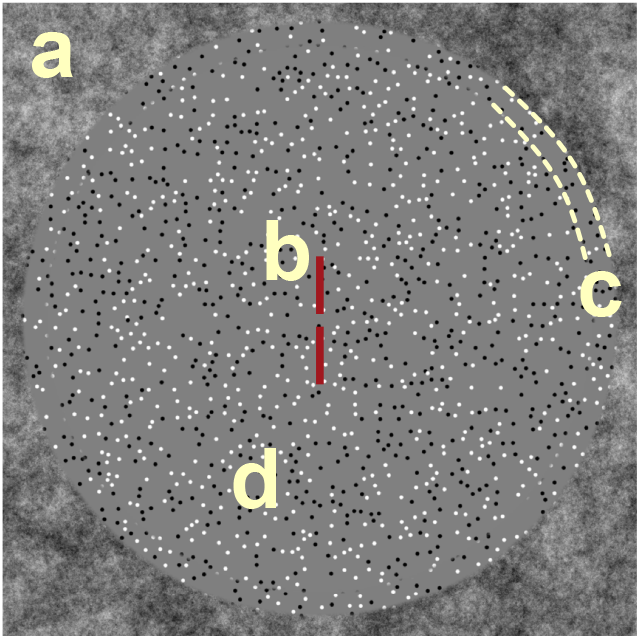
Schematic illustration of the DRDS display. A binocular, zero-disparity frame comprising 1/f noise (**a**) was used as a fusion lock. Dichoptic nonius lines (**b**) were used to engage fixation at the center of the screen and to monitor vergence. A detection task in which the color of the nonius lines changed from red to blue occurred at random intervals during both ‘attend fixation’ and ‘attend stimulus’ conditions. It was task relevant only in the ‘attend fixation’ condition. The duration of the color change was varied on a staircase which maintained an 82% detection rate. To minimize the availability of disparity references, a 1-degree gap containing binocularly uncorrelated dots (**c**) was interposed between the changing disparity region (**d**) and the fusion lock (**a**).

### Task and procedure

Two task conditions (Figure 1c) were tested on each of the two stimulus types. In the first condition, participants were instructed to press a button when a color change of the nonius lines occurred and were instructed to solely attend to this central fixation task (“ignore the dots”). The duration of the color change was varied (between 0.5 – 0.01 ms) using a staircase that maintained an 82% correct detection rate. Behavioral responses were tracked using the left arrow key on a keyboard. The DRDS stimuli were task-irrelevant in this condition. In the second condition, participants were instructed to attend to the DRDS stimulus while maintaining central fixation at the nonius lines (“fixate centrally but spread your attention to the dots”) and were asked to ignore the ongoing color change of the nonius lines. The DRDS stimuli were thus task-relevant in this condition. Each task condition was tested with both disparity plane and grating displays for a total of four experimental conditions (plane stimulus, attend fixation; plane stimulus, attend stimulus; grating stimulus, attend fixation; grating stimulus, attend stimulus).

Each stimulus trial lasted 12 seconds and was composed of a 1 s dynamic prelude to allow the EEG to achieve steady-state, followed seamlessly by a 10 s active task block, and ending with a 1 s dynamic postlude. The prelude and postlude recycled the first and last 60 frames of the stimulus presented during the active task block, respectively. Participants were instructed to blink as needed during jittered inter-trial intervals (1500 ± 500 ms). Each of the four conditions were presented in sets of 10 trials. Each condition set was repeated 3 times for a total of 30 trials per condition. The order of the sets was randomized and participants were permitted to rest between sets.

### EEG Acquisition and Pre-Processing

The EEG was recorded with 128-channel HydroCell sensor nets and an Electrical Geodesics Net Amps 400 amplifier (Electrical Geodesics, Inc., Eugene, OR, USA). EEG data were first bandpass filtered from 0.3 to 50 Hz, then evaluated according to a sample-by-sample thresholding procedure to identify consistently noisy individual sensors. These channels were interpolated by six neighboring channels. Once noisy sensors were substituted, the EEG was re-referenced from the Cz reference to the average of all 128 electrodes. Finally, 1 second EEG epochs containing a large percentage of data samples exceeding threshold (30 to 80 mV) were excluded on a sensor-by-sensor basis. These epochs were typically associated with eye movements or blinks. Independent components analysis was used on an individual participant basis (N = 8 of 20) to manually remove a systematic artifact arising on electrodes with higher 60 Hz power line noise. This artifact was characterized by sharp, clearly non-biological deflections of the EEG signal at regular intervals associated with the digital input trigger and were identifiable based on a characteristic topography, waveform, and Fourier spectrum.

### VEP Signal Processing and Reliable Components Analysis

The first step in our signal processing pipeline (Figure 7a) was to separate responses to the 20 Hz dot update rate from responses to the 1.4 Hz disparity change. This was done using the Fourier transform as a filter. Panel a(i) shows the spectrum of the response to a disparity grating on the left and the corresponding time average response on the right, reconstructed via an inverse Fourier transform. Panel a(ii) isolates the dot update response at 20 Hz (1F2) and 40 Hz (2F2, not shown). Panel a(iii) removes the dot update response, leaving only the harmonics of the disparity response (nF1) that do not overlap with the dot update rate in the reconstruction on the right. We also used the Fourier transform to filter the data into odd (1F1, 3F1, 5F1, etc.) and even (2F1, 4F1, 6F1, etc.) harmonic components as these have been found to have different functional properties (Kaestner et al., in revision).

**Figure 7:**
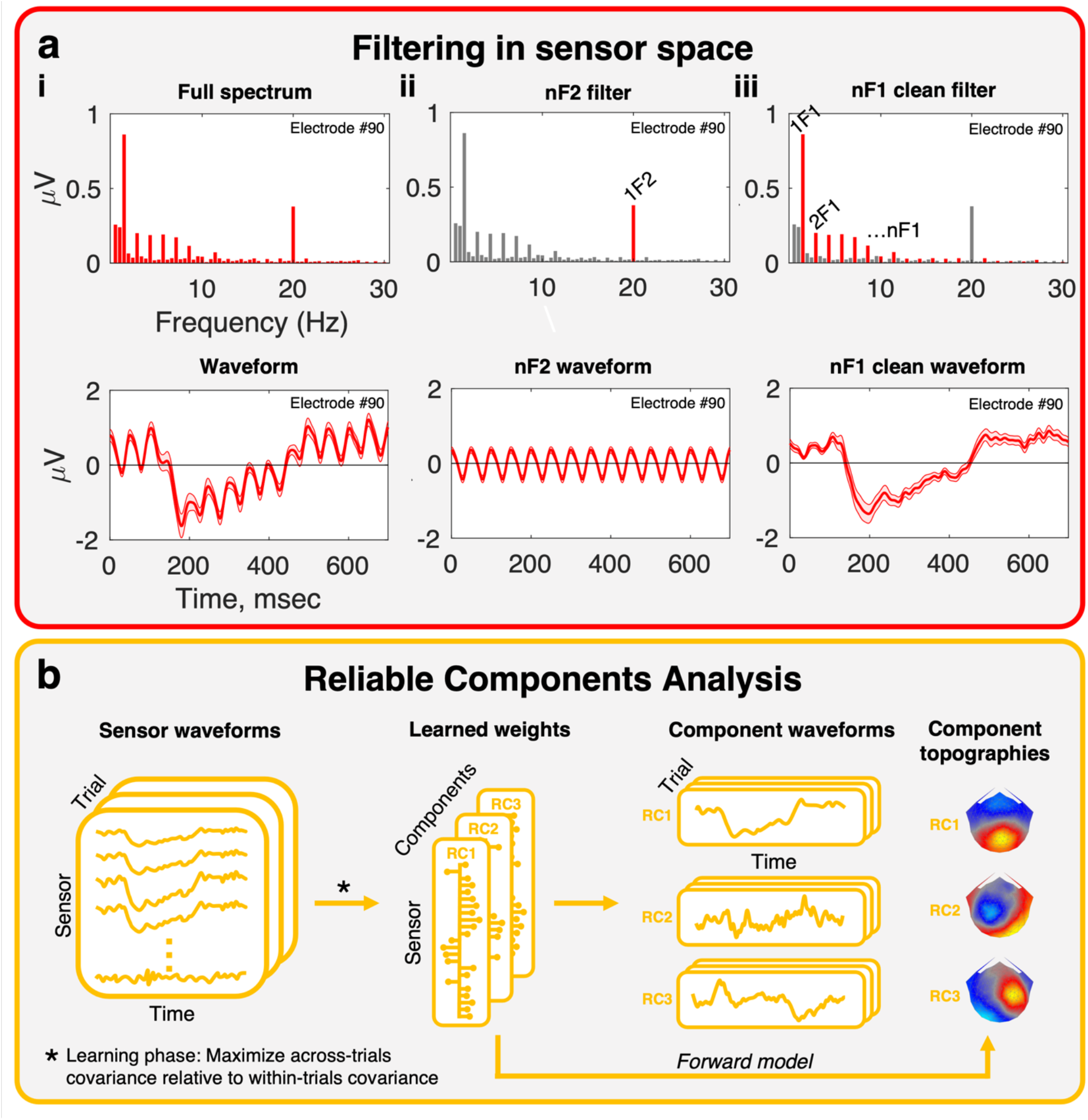
Two key stages in the data processing pipeline. **a:** Filtering in sensor-space. Fourier pairs (spectra on the top row, and corresponding waveforms on the bottom row) illustrate different filter types. The full spectrum, **a(i)**, contains both signal and noise harmonics, dominated by signals at multiples of the fundamental frequencies in the stimulus. This signal can be filtered to highlight different response components. The nF2 filter, **a(ii)**, removes all frequencies in the response except those at multiples of 1F2 (in our stimuli, the dot update rate at 20 Hz). The dot update can also be removed, and the disparity response highlighted, using the nF1 clean filter, **a(iii)**. Here, multiples of the 1F1 fundamental (in our stimuli, disparity update at 2Hz) are preserved and all other frequencies are removed. The reconstructed waveform is a “clean” version of the full waveform, with non-disparity and dot update signals removed. **b:** Dimensionality reduction via Reliable Components Analysis. The nF1 clean data are provided as input to the RCA pipeline, where components are learned that maximise the across-trials covariance relative to the within-trials covariance. Components are vectors of electrode weights where electrodes that respond in a consistent manner across trials are emphasized. These learned weights can be projected through a forward model, revealing underlying neural sources in the form of component topographies. Trial data are projected through the weight vectors to generate mean responses for each component, reducing the dimensionality of the data from 128 sensors to a small number of reliable components. Typically RCA is run on the group level, where input data are individual participant trials but weights are learned across all participants, resulting in group-level topographies and group-level waveforms.

The second step in the processing (Figure 7b) was to reduce the dimensionality of the data using Reliable Components Analysis (RCA) (Dmochowski et al, 2015). RCA is a spatial filtering approach akin to PCA, with the distinction that components are identified based on maximizing trial-by-trial covariance rather than variance explained ^55^. RCA computes linear combinations of electrode weights to maximize correlation, in time, of data across all trials of the same experimental condition. The optimization procedure, described in ^27,56^, reduces to an eigenvalue solution where the resulting weight vectors and their coefficients are the ordered eigenvectors and eigenvalues, respectively, of across-trial covariance relative to within-trial covariance ^57^. Thus, EEG data are decomposed from electrode-by-time matrices to component-by-time matrices, and components are sorted in descending order to maximize correlation in the first component. Finally, weight vectors are passed through a forward model ^27,28^ to reveal physiologically plausible topographies that indicate the neural sources likely to underlie the signal. For our purposes, we trained two sets of RCA filters on filtered data with the dot update response removed (‘nF1 clean’ filter), and components were learned across task conditions but within each disparity stimulus type. Our approach yielded two sets of weights representing the response to plane or grating type stimuli (forward models shown in Figure 2). In both cases the first reliable component, RC1, was dominant and subsequent components did not contain significant signals, thus we only present data from RC1.

### Permutation-Based Statistical Analysis

Timepoints where EEG responses from two different conditions differed from one another were localized using a permutation-based, paired-samples *t-*test which intrinsically corrects for multiple comparisons ^58-60^. Synthetic datasets were generated from the EEG data by randomly permuting condition labels across participants and calculating pairwise *t*-scores of the resulting waveform differences. For each permutation, we identified the longest consecutive run of time points where *p*-values were less than .050, generating a non-parametric reference distribution of significant consecutive timepoints across permutations. Consecutive trains of significant *t*-scores in the original, non-permuted data that exceeded 95% of the values in the reference distribution were considered statistically significant, and such epochs are marked with asterisks (Figures 3, 4 and 5). Since this procedure is dependent on both the length of the data and the arbitrary 95% threshold, we also present the uncorrected significance values (heat bars in Figures 3, 4 and 5).

## Acknowledgements

Funded by the National Eye Institute, National Institutes of Health, grant no. EY018875 awarded to AMN.

## Author contributions

MK and AMN designed the study. MK and YC collected and analyzed the data. MK produced the figures. All authors contributed to the writing and review of the main manuscript.

## Data availability statement

Data and code used for analysis will be made available on the OSF open science framework.

